# A unified derivative-like dopaminergic computation across valences

**DOI:** 10.1101/2025.08.11.669800

**Authors:** Mijoo Park, Jongwon Yun, Hogyu Choi, HyungGoo R. Kim

**Author notes:** Correspondence: HyungGoo R. Kim.

## Abstract

Dopamine activity in the brain affects decision-making and adaptive behaviors. A wealth of studies indicate that dopamine activity encodes discrepancy between actual and predicted reward, leading to the reward prediction error (RPE) hypothesis. Specifically, it has been claimed that mesolimbic dopamine activity conforms to temporal-difference reward prediction error (TD RPE), a teaching signal in machine learning algorithms. Recently, there is growing evidence suggesting that dopamine is also involved in learning during aversive situations. However, the fundamental computation of dopamine activity in aversive situations is still unknown. A plausible but untested hypothesis is that dopamine activity in aversive situations also encodes TD RPE. Here, we tested this hypothesis by using mice in virtual reality. Mice were trained to avoid electrical tail shocks by running out of a virtual shock zone. Using probe conditions with speed manipulation or teleportation, we revealed that the dopamine signal in the ventral striatum follows the temporal derivative form of a value function. Delivering a reward at the end of the track enabled us to observe the integration of aversion and reward in a derivative form. Moreover, the value functions unbiasedly estimated from the recorded signal is consistent with the initial hypothetical form, with a realistic reflection of a received shock distribution. Taken together, our results show that mesolimbic dopamine activity can operate as a unified teaching signal in natural situations with positive and negative valences.

## Introduction

Living organisms adaptively change behaviors to maximize reward and minimize punishment. Dopamine (DA) has long been recognized to be a critical neuromodulator in adaptive processes such as learning, motivation, and movement (Bromberg-Martin & Hikosaka, 2009; Schultz, 2002; Wise, 2004). Numerous studies have shown that dopamine neurons respond to reward and sensory cues that predict reward (Eshel et al., 2015; Glimcher, 2011; Schultz, 2002; Schultz et al., 1997; Watabe-Uchida et al., 2017), and the manipulation of DA activity reinforces behaviors (Steinberg et al., 2013). A combination of theoretical and experimental work demonstrated that dopamine activity conforms to temporal-difference reward prediction error (TD RPE) in reinforcement learning, a ‘teaching signal’ that changes value memory and behaviors (Kim et al., 2020; Niv, 2009; Schultz et al., 1997). TD RPE (δ_*t*_) is defined as:

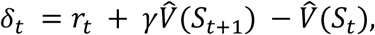

where *r*_*t*_ is the reward given to an animal at time *t*, *S*_*t*_ is the state that animal is in at time *t*, and γ works as a discounting factor for future rewards. *V̂* (*S*_*t*_) is the sum of discounted future rewards in the state at time *t* (Sutton & Barto, 1998). If immediate reward or punishment is absent (*r*_*t*_ is zero) and γ is close to 1, TD RPE is approximately the derivative, or slope, of value function, *V̂*(*S*_*t*_). Based on this relationship, it has recently shown that slowly fluctuating dopamine activity associated with goal approach (Howe et al., 2013) also represents a moment-by-moment TD RPE (Kim et al., 2020; Mikhael et al., 2022).

While these studies elucidated the nature of adaptive processes mediated by dopamine in ‘good’ situations, whether and how dopamine is involved in decision-making in negative situations is still unclear. However, recent work has shed light on the role of dopamine in aversive situations (Lopez & Lerner, 2025). As an example, VTA dopamine neurons are inhibited by aversive stimuli (H. Matsumoto et al., 2016; M. Matsumoto & Hikosaka, 2009) and, in classical conditioning tasks, dopamine activity in the NAc core is inhibited by cues that predict an aversive stimulus (Badrinarayan et al., 2012; de Jong et al., 2019; Oleson et al., 2012; Stelly et al., 2019). Beyond the Pavlovian conditioning paradigms, increased DA responses were shown during successful avoidance of negative stimuli like foot shocks (Lopez et al., 2024; Oleson et al., 2012; Stelly et al., 2019), suggesting that avoidance behavior can be reinforced similarly to reward-based learning. In well-trained subjects, while DA is inhibited to aversive cues, DA is increased during safety periods, and the excitation is correlated with successful avoidance (Kutlu et al., 2021; Stelly et al., 2019). Furthermore, optogenetic stimulation of DA neurons during the conditioned stimulus (CS) period led to improved avoidance behavior (Wenzel et al., 2018). These results suggest that dopamine is involved in adaptive behaviors in aversive situations, but the neural mechanisms are unknown.

An important open question is whether dopamine neurons compute moment-by-moment TD RPE in aversive situations, as in rewarding situations. This idea assumes that once animals recognize an aversive area, the value of the place would be lower than the baseline, and dopamine activity still encodes the derivative of the value function. To test this, we designed an active avoidance task in mice using virtual reality (VR). The virtual track included a region in the middle where mice could potentially receive electric tail shocks. Once mice learned that the middle position was aversive, we investigated the nature of dopamine activity by adding ‘probe’ conditions, such as manipulating animal’s position or speed gain. We show that dopaminergic activity in the ventral striatum during aversive situations also conforms to TD errors. Our results suggest that dopamine signals follow the derivative form of value function in both positive and negative valences, further explaining the role of dopamine in learning and decision making.

## Results

### Dopamine activity shows ramp down followed by excitation during an avoidance task in virtual reality

To create an environment where an animal encounters and avoids aversive situations, we positioned a ’shock zone’ in the middle of a virtual linear track (Fig. 1A, Video 1). In the shock zone (60 a.u. – 160 a.u., 1 a.u. = 0.95 cm), mice received mild electric tail shocks (Fig. 1D; 0.8 mA, 150 ms duration, 3.5 s interval) if they stayed stationary (running speed < 4 a.u./s). The tail shock caused mice to exhibit transient running behavior at high speed (Supp. Fig. 1B, n = 14 mice). Initially, mice largely failed to avoid this zone and received multiple electric shocks. Across training days, the proportion of failed trials, in which mice received at least one shock, decreased markedly (Supp. Fig. 1A, R = -0.97, P = 0.0002, Spearman correlation, n = 14 mice). Once mice learned that the middle zone could potentially give shocks, they stopped voluntarily walking toward the shock zone. Therefore, we made the visual scene passively progress (minimum speed of 2 a.u./s) if mice did not move only in the beginning zone (see Methods). As mice learned the task, we observed that mice initiated running right before entering the shock zone and stopped right after they exited the shock zone (Fig. 1E). We argue that this anticipatory running at the entrance of the shock zone is the behavioral evidence that mice associated the shock zone with aversive stimuli, analogous to anticipatory licking in rewarding situations (Cohen et al., 2012; Kim et al., 2020). The speed of anticipatory running (Fig. 1F) was significantly greater than the baseline speed (Fig. 1G, P = 0.0005, signed-rank test, n = 14 mice; P < 0.01 for all individual sessions) and it was higher in the last day of training than the first day (Supp. Fig. 1C, P < 0.05, signed-rank test, n = 14 mice).

**Figure 1.**
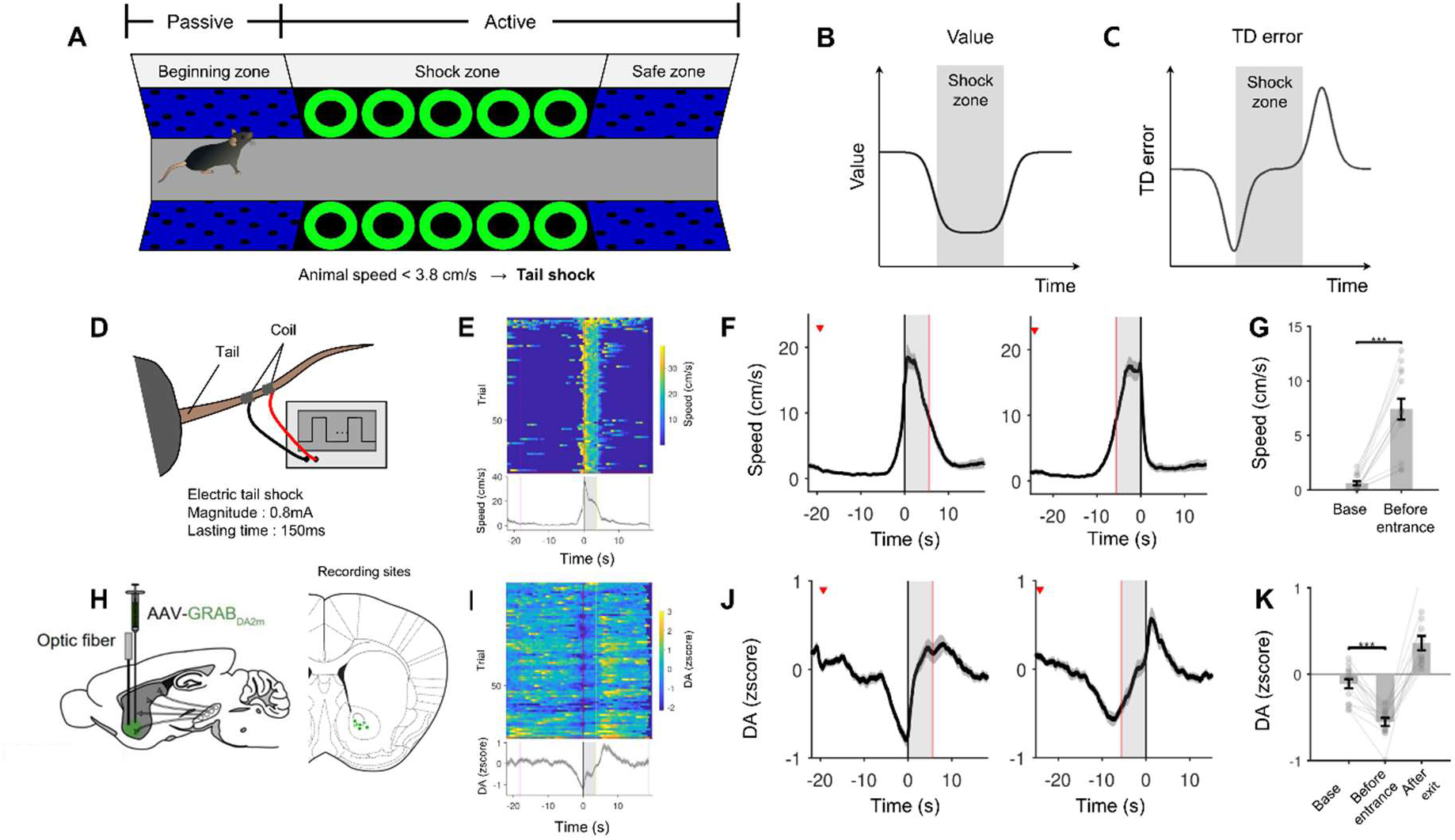
Behaviors and dopamine activity in the standard avoidance task. (A) The virtual linear track in the standard experiments (Track 1). (B) State value over position (Kim et al., 2020). Shaded range depicts the shock zone. (C) Temporal difference error predicted from the state value assuming constant speed. (D) Schematics of the tail of the mouse and the coil to deliver electrical shocks. (E) Locomotion speed from an example session. Individual trial results are aligned by the shock zone entrance (top) and averaged (bottom). (F) Average speed aligned by the time of shock zone entrance (left, n = 14 mice) and shock zone exit (right, n = 14 mice). Red line indicates the median shock zone exit (left) and shock zone entrance (right), respectively. (G) Comparisons of speed between baseline -12s to -10s)and anticipatory running right before shock zone entrance (-3s to 0s). (H) Surgery for fiber photometry. (I) Dopamine activity of the example session (E) Dopamine activity aligned by the shock zone entrance. (J) Average dopamine activity of aligned by the time of shock zone entrance (left, n = 14 mice) and shock zone exit (right, n = 14 mice). Red line means the median shock zone exit (left) and shock zone entrance (right), respectively. (K) Comparisons of dopamine activity between baseline [-12s -10s], right before entrance [-5s 0s], and shock zone exit, [0s 5s]. Error bars and shadings indicate s.e.m., one, two, and three stars indicate P values less than 0.05, 0.01, 0.001, respectively.

Once mice learn the task, the value for the shock zone would be lower than the two other zones (Fig. 1B). If the dopamine signal reflects moment-by-moment TD error also in this negative situation, we would observe a decrease of dopamine signal in the entrance of the shock zone and the excitation at the exit of the shock zone (Fig. 1C). We injected a GRAB-based dopamine sensor (DA2m, Sun et al., 2020) and recorded dopamine activity in the ventral striatum (targeting nucleus accumbens core) using fiber photometry (Fig. 1H, see Methods). As mice approached the shock zone, dopamine activity gradually ramped down until animals reached the shock zone, which was followed by an excitation as the mice ran out of the shock zone (Fig. 1I). These neural trends were consistent across populations (Fig. 1J). Additionally, the dopamine signal was significantly lower before entrance (Fig. 1K, P = 0.0001, signed-rank test, n = 14 mice) and higher after exit (Fig. 1K, P = 0.0017, signed-rank test, n = 14 mice) compared with the baseline signal, which conformed to the TD error prediction (Fig. 1C). The inhibition of dopamine signal was rapidly developed on the first day of training as the animals experienced shocks (Supp. Fig. 2A) and this ramp-down activity was more clearly shown in later trials compared with the initial trials (Supp. Fig. 2B, P < 0.0001, R = -0.25, Spearman correlation, n = 14 mice). Aligning the same data at the exit of the shock zone shows gradual and rapid formation of excitation on the first day (Supp. Fig. 2C-D, P = 0.0002, R = 0.14, Spearman correlation after excluding the first two trials, n = 14 mice).

Dopamine response dropped down at the visual stimulus onset once mice learned the task (Supp. Fig. 2E-F, P = 0.0007, signed rank test, n = 14 mice), and increased after stimulus offset compared with the signal before visual stimulus offset (Supp. Fig. 2G-H, P = 0.012, signed rank test, n = 14 mice). Dopamine response to the shock varied across individuals; some animals showed phasic excitation, while others showed more flat response in both the first day and last day of training (Supp. Fig. 1D, n = 14 mice).

### Speed gain modulates dopamine activity in both negative and positive directions

Upon the completion of the standard avoidance task, we further examined the nature of dopamine activity by performing additional experiments with ‘probe’ conditions. First, we added conditions with variable speed gain for the visual scene. In the new conditions, visual stimuli moved at twice or half as fast as in the standard condition. If the animal learned a value function over position and dopamine activity is the derivative (slope) of the value function (Fig. 2A), the magnitude of both ramp-down and excitation would increase with speed (Fig. 2B). However, if for instance dopamine response directly reflects the state value itself, the range of inhibition and excitation should be minimally impacted by scene speed.

**Figure 2.**
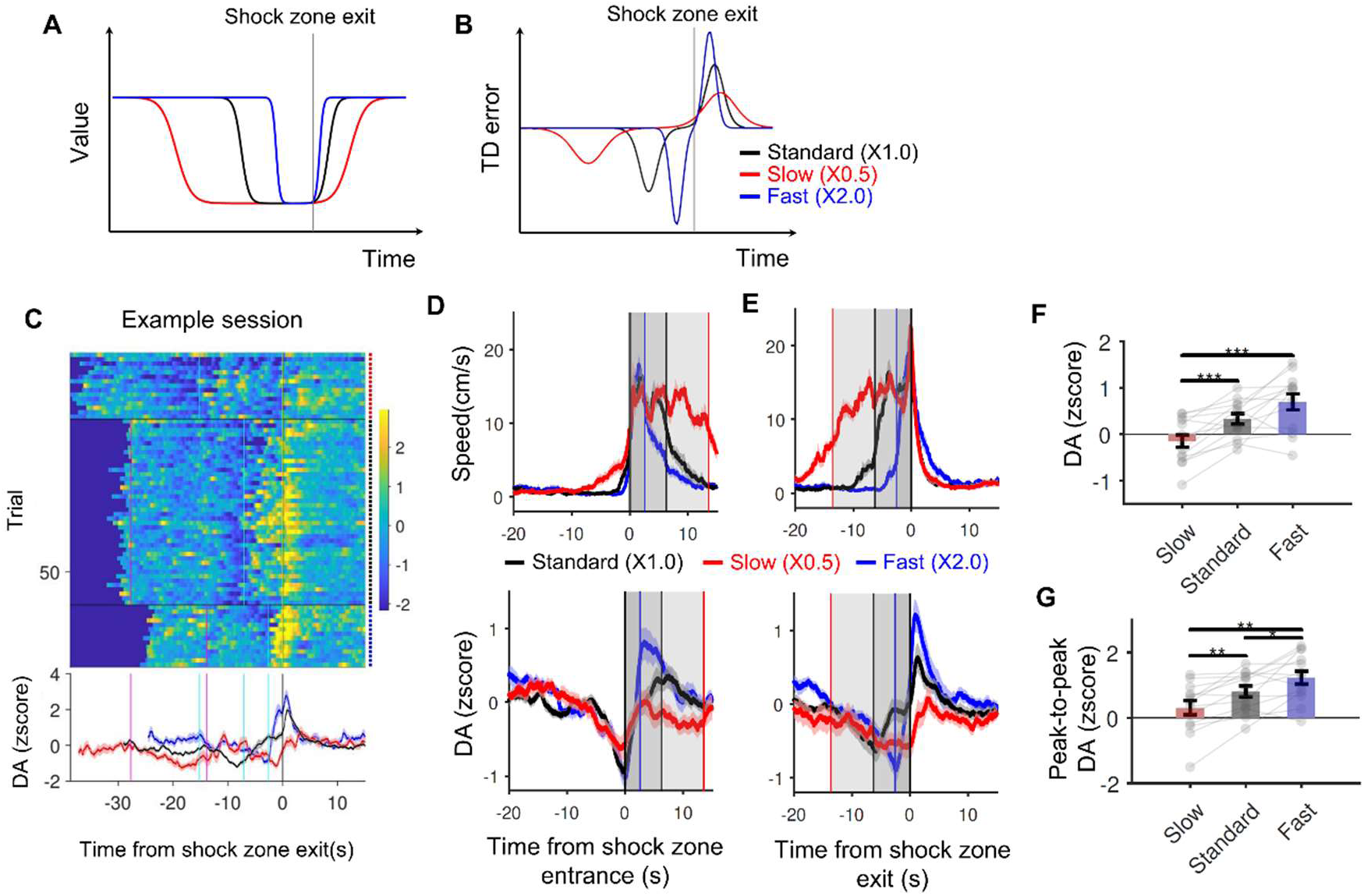
Speed gain manipulation experiment. (A) State value as a function of time in difference speed gain conditions aligned by shock zone exit. (B) TD error predicted from the state value (A). (C) Example session for speed gain experiment. (D) Average speed aligned by the time of shock zone entrance (left, n = 13 mice) and shock zone exit (right, n = 13 mice). Lines mean the median event time of shock zone exit (left) and shock zone entrance (right) for each color-matching condition. (E) Average dopamine signals aligned by the time of shock zone entrance (left, n = 13 mice) and shock zone exit (right, n = 13 mice). Lines mean the median event time of shock zone exit (left) and shock zone entrance (right) for each color-matching condition. (F) Comparisons of dopamine signal of different conditions by time window of [0s 5s] from shock zone exit (P = 0.0007 (slow and standard), 0.0007 (slow and fast), signed rank test, n = 13 mice). (G) Comparisons of dopamine signal of subtracting quantified data of [-5s 0s] in shock zone entrance from quantified data in time window of [0s 5s] from shock zone exit (P = 0.0012 (slow and standard), 0.0012 (slow and fast), and 0.0269 (standard and fast), signed rank test, n = 13 mice).

We randomly interleaved the visual gain of 0.5, 1, and 2 (see Methods; Fig. 2B, Video 4). If the aversiveness of the shock zone is static, the TD error model predicts that both the magnitudes of ramp-down and excitation would increase with speed gain as the slope of the value function depends on the scene speed in the virtual track.

As expected, the running duration of animals decreases with speed gain (Fig. 2C). We observed a greater DA ramp-down and excitation (Fig. 2C) in the fast speed condition, compared with the slow speed condition. The trend was consistent in the population data (Fig. 2E bottom and 2F, R = 0.57, P = 0.0002, Spearman correlation between three levels of speed and DA response, n = 13 mice), and the peak-to-peak amplitude of the dopamine signal significantly increased with the speed gain (Fig. 2G, R = 0.51, P = 0.0008, Spearman correlation between three levels of speed and DA response, n = 13 mice). These results were consistent with the prediction that dopamine activity reflects the temporal derivative of a value function in aversive situations. Note that the magnitudes of ramp-down were not significantly different between the standard and fast conditions (Fig. 2F, P = 0.0574, signed-rank test, n = 13 mice). We reason that this may be attributed to a floor effect, due to the low firing rate of dopamine neurons.

### Dopamine response to teleportation is explained by an abrupt change in the value function

We further investigated dopamine activity using a sudden change in position (teleportation). If mice estimate value based on the position, teleportation from the shock zone entrance to the exit (Fig. 3A-B, red arrows) would lead to a sudden increase in value, which may produce transient excitation (Fig. 3C, red). Using a new track with different patterns between initial and safe zones (Track 2, Fig. 3A), the mouse was teleported to skip the shock zone (Fig. 3A, Video 5). Since teleportation incorporates a sudden change of the visual pattern, the transient dopamine responses to such a change of visual scene have the potential to be explained by sensory salience or sensory prediction error (Stalnaker et al., 2019; Takahashi et al., 2017). Thus, we added a backward teleport condition, changing the scene back to the entrance of the shock zone again right after the animal passed the exit (Fig. 3A-B, blue arrows). If the response reflects sensory surprise, we will also observe excitation after backward teleportation. However, if the dopamine signal encodes TD error, a transient inhibition would follow after backward teleport (Fig. 3C, blue) caused by the sudden decrease of the state value.

**Figure 3.**
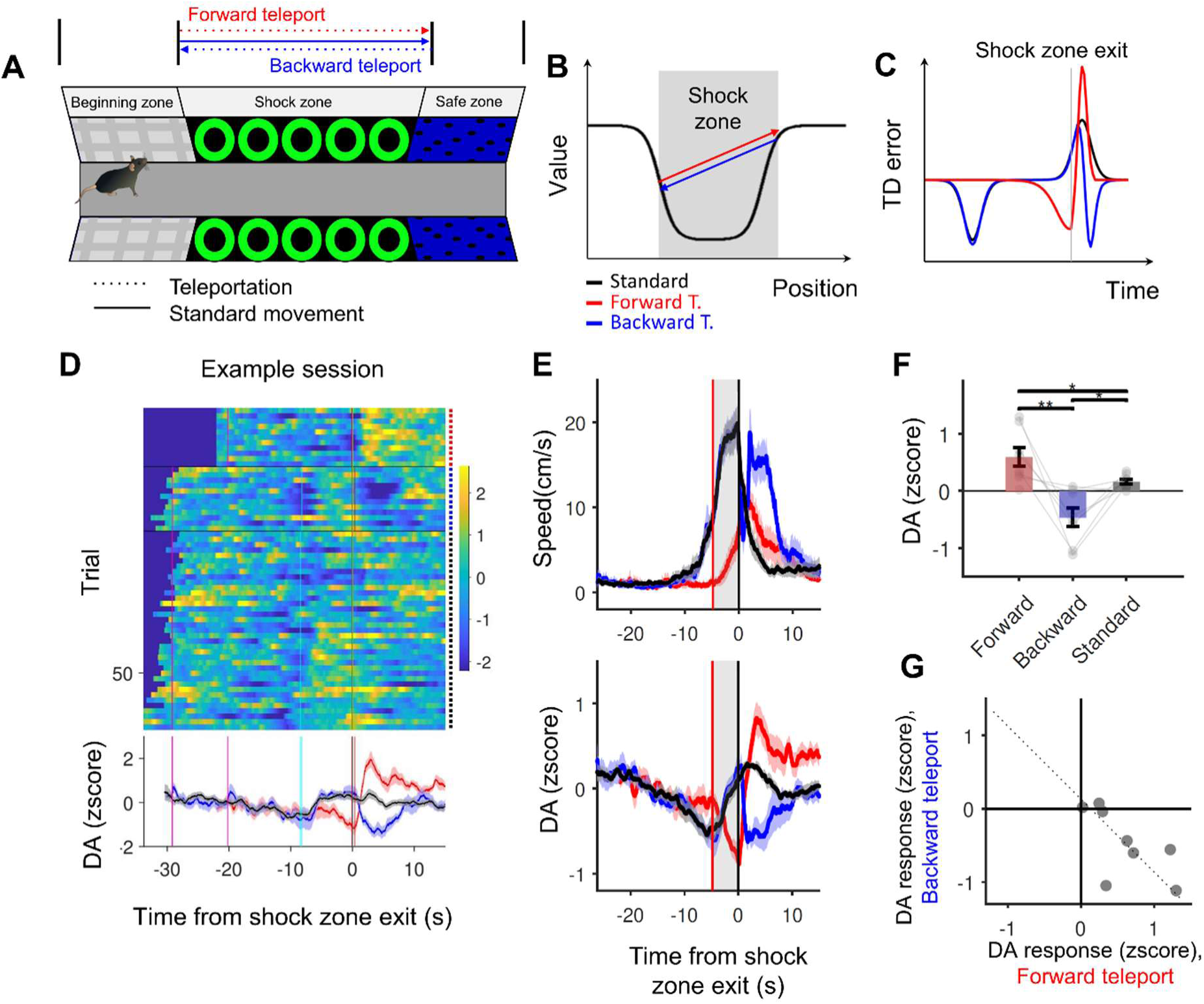
Teleportation experiment. (A) The virtual linear track type 2. (B) State value in function of time. Shaded windows can be the time when mice are inside the shock zone. (C) Temporal difference error predicted from the state value (A) aligned by teleport event. (D) Example session for teleport experiment. (E) a Population result of Locomotion (speed) result each aligned by the time of shock zone exit (n = 8 mice) for each color-matching condition. (F) Population result of dopamine signals each aligned by the time of shock zone exit (n = 8 mice). Red line means the average shock zone entrance timing. (G) Comparisons of dopamine signal of different conditions by time window from shock zone exit, [0s 5s] after the shock zone exit (standard, black) and teleport (red and blue) (P = 0.0078 (forward and backward), 0.0234 (backward and standard), and 0.0156 (standard and forward), signed rank test, n = 8 mice). (H) Correlation between quantified forward teleport response and backward teleport response (P = 0.022, r = -0.81, Spearman correlation, n = 8 mice).

Our results showed significant excitation in response to the forward teleportation and significant inhibition after backward teleportation (Fig. 3D), supporting the idea that dopamine response follows moment-by-moment derivative of value function in aversive situations. This was consistent in population results (Fig. 3E bottom). Phasic excitation after forward teleport is significantly greater than the response in the standard condition (Fig. 3F red and gray, P = 0.0156, signed-rank test compared with standard condition, n = 8 mice), whereas the inhibition after backward teleport is significantly smaller than the response in the standard condition (Fig. 3F blue and gray, P = 0.0234, signed-rank test compared with standard condition, n = 8 mice). However, although the modulation is significant on average, some animals did not exhibit robust modulation of dopamine signal in the teleport conditions (3 mice out of 8 mice). We reasoned that the variability in the shape of the value function may underlie these results. For example, if a mouse’s value function begins to drop behind the shock zone entrance, the same forward teleport may not induce large change in value in these animals (Supp. Fig. 3B). Importantly, in these animals, backward teleportation would also induce small inhibition. Indeed, this was exactly what we observed. There was a significant negative correlation between the transient excitation and inhibition of dopamine signal caused by forward and backward teleport (Fig. 3G, P = 0.022, R = -0.81, Spearman correlation, n = 8 mice). We further tested this possibility by direct manipulation of teleport begin and end positions. The two mice initially did not show significant excitation or inhibition of dopamine (Supp. Fig. 3C) with the standard teleport start and end positions (Supp. Fig. 3A middle). However, with a slight shift in the teleport positions (Supp. Fig. 3A bottom), We observed significant excitation and inhibition in both animals (Supp. Fig. 3D).

### Derivative-like dopamine activity integrates aversive and rewarding outcomes

So far, we have measured dopamine activity during aversive situations without actual rewards. In natural situations, animals often pass through dangerous areas with predators to obtain food or water. We wondered whether dopamine activity can provide a derivative-like signal in more complex situations. We added a new Reward Track (Track 3, Fig 4A, Video 6) in which water-deprived mice obtain a drop of water (0.3ul) 1.5s after they exit the shock zone. Unlike track 1 without reward, reward would make the value in the safe zone greater than that of the beginning zone, resulting in greater dopamine response. We randomly interleaved the Reward (50%) and Standard (50%) Tracks.

**Figure 4.**
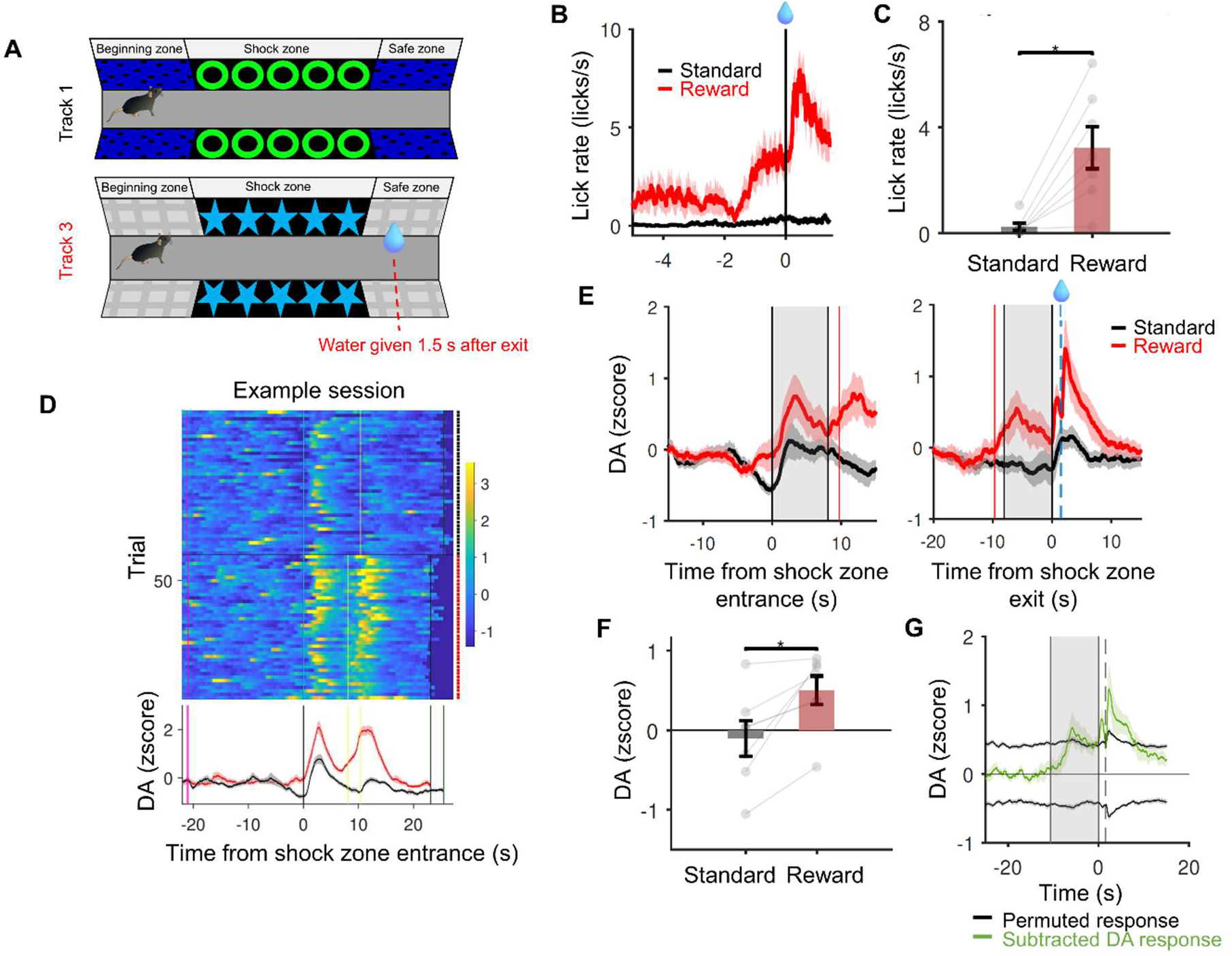
Integration of aversion and reward. (A) The virtual linear track 3 including reward compared with track 1 used in standard condition. (B) Population result of lick data aligned by the time of reward onset (n = 7 mice) for each color-matching condition. (C) Comparison of lick results quantified by the time window of [-2s 0s] from reward onset (P = 0.016, signed rank test, n = 7 mice). (D) Example session of reward added experiment. (D) Dopamine signals of a population result are each aligned by the time of shock zone entrance (n = 7 mice). Lines mean the average shock zone exit timing for each color-matching condition. (E) Dopamine signals of a population result are each aligned by the time of shock zone exit (n = 7 mice). Lines mean the average shock zone entrance timing for each color-matching condition. (F) Comparisons of dopamine signal of different conditions by time window from shock zone exit, [0s 1.5s] right before the reward is delivered (P = 0.013, signed rank test, n = 7 mice). (G) Subtraction of standard condition dopamine response from reward condition dopamine response (green, n = 7 mice). Black lines are 5% and 95% confidence intervals computed by permutation test (see Methods, n =7 mice)

Mice showed anticipatory licking only in Track 3 (Fig. 4B-C, P = 0.016, signed rank test, n = 7 mice), indicating that they successfully learned the existence of reward in Track 3 with different visual cues. If the value is high at the end, the increase of dopamine signal would be greater as mice get out of the shock zone. Dopamine signals in the reward-given track show greater dopaminergic activity before getting reward than those in the standard condition (Fig. 4D), and this effect generalized well in population data (Fig. 4E-F, P = 0.013, signed-rank test, n = 7 mice). To estimate the net effect of reward, we subtracted the response in the standard condition from the response in the reward-given condition (Fig. 4G, green). The net response right before reward is significantly greater than the baseline (Fig. 4G, permutation test, see Methods), such that the differential dopamine activity results in a ‘ramp-up’ right before reward delivery as shown in previous work (Kim et al., 2020).

### Model-fit confirms the derivative nature of DA activity and visualizes value function

The design and interpretation of our results assumes that the value of the shock zone would be lower than the other zones (Fig. 1B). Here, we examined the nature of dopamine activity without such an assumption. Instead, we adopted a model-fit procedure to estimate the shape of the value function, defined by 10th-degree polynomial function. Predicted dopamine activity was computed from the value function and position time courses. The model-fit procedure iteratively compared this predicted dopamine activity and empirical data until it found the best-fit parameters for the value function and discount factor (see Methods, Fig. 5A, green). An example result shows that the TD model prediction followed the trend of recorded data (Fig. 5A). We also tested an alternative model that dopamine activity encodes value itself, not the temporal derivative of value function (Hamid et al., 2016). In this case, the model-fit procedure attempted to find the best-fit parameters of the value function that matches measured dopamine activity (Fig. 5A, purple). For each animal, average responses from available standard, speed, and teleport experiments were combined to increase statistical power of model comparisons.

**Figure 5.**
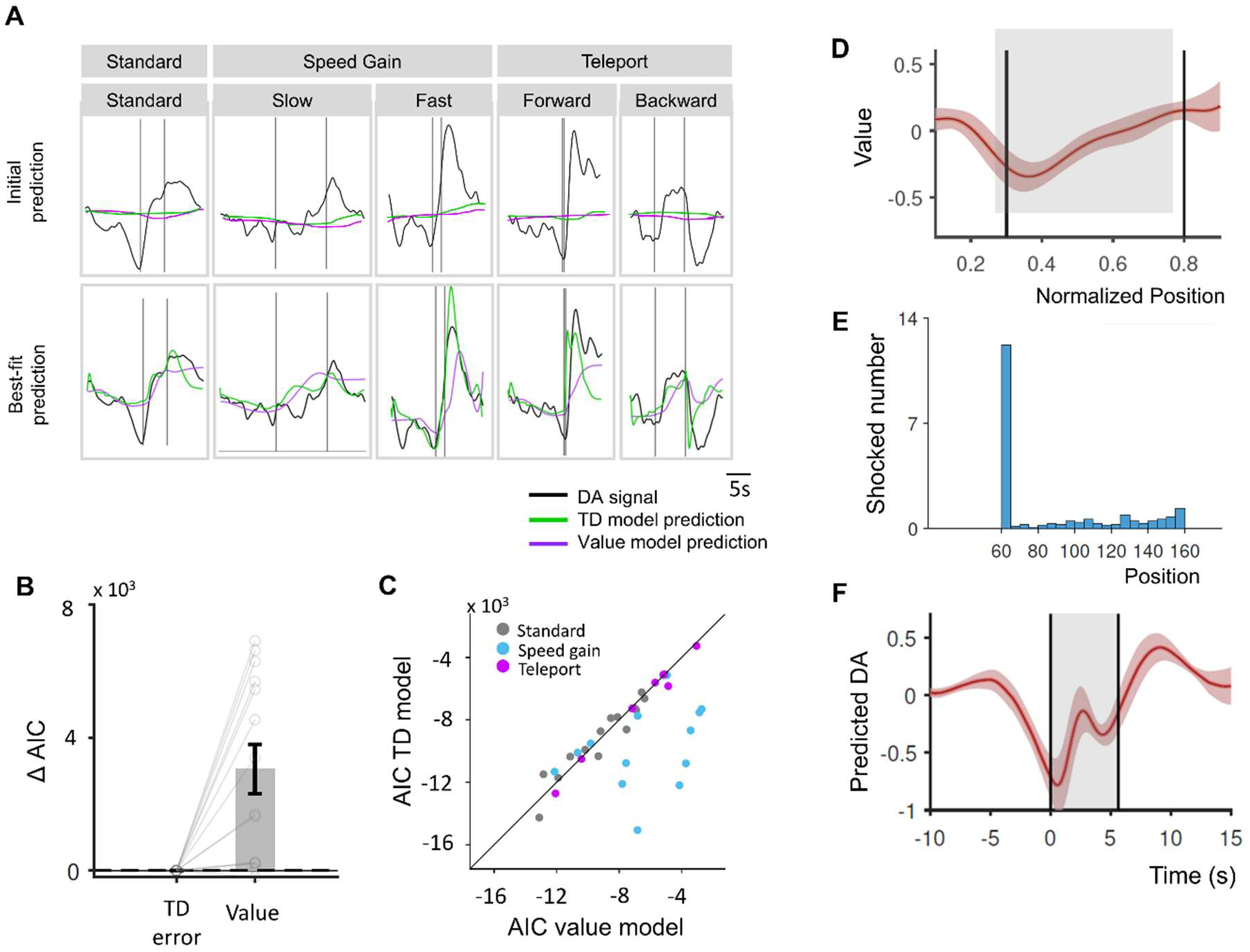
Model comparisons and recovered value function. (A) Best fitted prediction (green) from model fitting and recorded dopamine signal (black) (B) Comparison of AIC between TD error model and value model (P = 0.0004, signed rank test, n = 14 mice). (C) Correlation between AIC of the value model and TD model. Each dot represents each session. (D) Recovered value function from fitted parameters. (E) Average shock number in a session at the last day of training standard condition. (F) Predicted dopamine response with DA2m filters.

We used Akaike information criterion (AIC) to compare model-fit performances. AIC was lower in the TD model, implying that the TD model can explain the data better (Fig. 5B, P = 0.0004, signed-rank, n = 14 mice). Although model performances using data in the Standard Track did not differ significantly, using data in the probe experiments resulted in lower AICs for the TD model, especially in the speed gain manipulation (Fig. 5C), which means that the fitting process itself is not biased, but results from manipulation experiments are necessary to manifest the temporal difference computation in dopamine activity.

The recovered value function from the best fit parameters (Fig. 5D) resembles our schematic illustration (Fig. 1B), but with some differences in detail. The value function shows a trough at the entrance followed by increase within the shock zone. We reasoned that the discrepancy may be attributed to the empirical position where the mice were punished. Indeed, mice received many more shocks as they entered the shock zone (Fig. 5E). The predicted dopamine activity in the standard condition (Fig. 5F) resembles our measured data (Fig. 1J).

## Discussion

Previous studies investigating the computational nature of dopamine activity have mainly focused on rewarding situations. However, recent evidence suggests that dopamine is also involved in processing aversive situations. Here we examined whether mesolimbic dopamine signaling is a moment-by-moment derivative of value function in aversive situations. Using mice that associated a specific zone with aversive events in virtual reality, we tested the central tenet of TD RPE by manipulating the position of animals in space. Our results unambiguously demonstrate that dopamine activity in the ventral striatum encodes moment-by-moment TD error in aversive situations. These results were further verified by unbiased model fitting that did not assume a specific value function. Finally, we have shown that dopamine signals integrate both rewarding and aversive information, showing that dopamine activity can be used as a unified teaching signal across valences.

### A unified derivative computation for learning and decision-making extending TD RPE framework to aversive learning

It has been a fundamental question in neuroscience to understand how humans and animals adaptively change behaviors. Reinforcement of behaviors by reward and punishment have been two major sources of behavioral adaptation. Numerous experiments have shown that dopamine is a key neuromodulator in reward-based learning, and the TD RPE hypothesis has provided one of the most successful theoretical accounts to explain both neural activities and behaviors. Leveraging the slow fluctuations during approach-to-goal tasks, we previously showed that dopamine neurons compute TD RPE in a moment-by-moment manner (Kim et al., 2020).

Here, we asked whether the same computational principle can be applied while animals confront aversive situations. This is not an obvious question, given that different peripheral afferents and brain areas provide inputs for positive (e.g., water) and negative (e.g., electrical shock) valences. Appetitive reward signals are provided by hypothalamus, whereas brain areas such as periaqueductal gray (PAG) and insula cortex are involved in punishment. Previous studies show that some mesolimbic dopamine neurons are inhibited by punishments and cues predicting punishment. Our work extended the previous study to the level of neural computation (Bromberg-Martin et al., 2010; Cox & Witten, 2019) - dopamine neurons compute the unified derivative computation between positive and negative valences.

### Relationships with previous studies regarding dopamine activity in aversive situations

Dopamine activity in our task shows ramp-down right before the shock zone entrance followed by modest excitation. These patterns are somewhat different from previous studies using fear conditioning (Badrinarayan et al., 2012; de Jong et al., 2019; Oleson et al., 2012) or avoidance paradigms. Most studies showed transient inhibition at the time of cue onset, and transient excitation at the cue indicating no punishment (Kutlu et al., 2021; Stelly et al., 2019).

Why do we observe activity patterns that are qualitatively different from previous studies? We reason that how the sensory cue was presented contributes to the difference. Previous studies have mostly used discrete tone or visual stimulus as a cue (Badrinarayan et al., 2012; de Jong et al., 2019; Oleson et al., 2012). In our task, however, animals witnessed that the shock-zone pattern gradually approached them. As a result, mice observed dynamic sensory evidence that suggested proximity of potential punishment. We have previously shown that dynamic sensory evidence may induce dopamine ramp up in rewarding situations (Kim et al., 2020). We believe that the same mechanism but with opposite valence may underlie the dopamine ramp down in our current experiments. Similarly, as the animal left the shock zone, they observed the shock zone pattern gradually disappeared in their sight. We believe that it is why the excitation is not highly transient. In contrast, in the forward teleport condition, the stepwise increase of value function from negative back to the baseline induces transient excitation in some animals (Fig. 3E bottom, red). We speculate that the ‘safety signal’ in previous studies can be largely explained by the stepwise change of value, with the dopamine activity being the derivative of the value function.

Other previous work has demonstrated that excitation of dopamine neurons in cue presentation can predict avoidance (Oleson et al., 2012), and the excitatory response to cue seems to be at odds with our results. In operant tasks where action reliably induces shock omission, it could be possible that the value function increases at the moment when animals decide to press the lever because under TD learning, positive prediction error can be propagated backward to the clear sensor or motor action (Schultz et al., 1997).

### Interpretation of results and remarks on alternative explanations

Dopamine is also known to be related to motivation (Salamone & Correa, 2012), vigor (Panigrahi et al., 2015) and movements (Howe & Dombeck, 2016). In our tasks, mice appear to be motivated to avoid the shock zone and execute locomotive movement. In the standard experiments, motivation to avoid would gradually increase as animals approach the shock zone, whereas the dopamine responses gradually decrease. During the significant ramp-down, mice did not show meaningful movements. Our probe conditions further provide evidence that these possibilities may not explain our results. In the speed gain conditions, mice show a similar range of speed but dopamine activity is markedly different. In teleport conditions, mice did not exhibit much running or sudden movements, but dopamine still showed clear transient excitation or inhibitions, consistent with the expected changes of value function. These manipulated conditions show that the increase and decrease of dopamine signal is not solely caused by movement.

Another possibility is that mice may consider the safe zone as rewarding (Stelly et al., 2019). In the reinforcement learning framework, it means that *r*_*t*_ in the TD error equation can be positive. The positive *r*_*t*_ may explain the excitation in the standard condition without dopamine response being the derivative signal. However, it cannot explain significantly larger excitation in the forward teleport condition (Fig. 3F red). This is why the probe conditions are critical in examining the nature of dopamine responses. Similarly, it cannot explain greater peak-to-peak measurements in the fast speed condition (Fig. 2G).

### Diversity of dopamine systems and future directions

While we focus on the mesolimbic dopamine pathway projecting to the ventral striatum, there is now mounting evidence that dopamine neurons encode distinct information such as emotional salience and threat prediction error, depending on the projection target (C. K. Kim et al., 2016; Menegas et al., 2017; Parker et al., 2016) or subtypes (Azcorra et al., 2023). For example, dopamine responses in the prefrontal cortex (Sorg & Kalivas, 1993; Yoshioka et al., 1996), dorsal striatum (Budygin et al., 2012; Ilango et al., 2012), tail of striatum (Tsutsui-Kimura et al., 2025; Zafiri et al., 2025), and amygdala (Tang et al., 2020; Young & Rees, 1998) appear to increase in response to aversive stimuli. Therefore, detailed characterization is required to further investigate the neural computation underlying these diverse dopamine responses. We believe that our experiments can provide valuable toolkits in these characterization processes, especially if combined with genetic tools that can dissect subtypes of dopamine populations.

Among the cortical and subcortical subregions, we believe that the comparison between ventral striatum and tail of striatum would be very intriguing. Previous study highlighted that tail of striatum (TS)-projecting dopamine neurons were activated in response to cues predicting aversive outcomes, such as shocks (Badrinarayan et al., 2012). Also, dopaminergic neurons projecting to the TS are not only involved in threat anticipation but are also particularly sensitive to novel and unexpected stimuli (Menegas et al., 2018), which elicit an approach-avoidance conflict (Tsutsui-Kimura et al., 2025). This sensitivity to novelty and threat-related cues implies that TS-projecting dopamine neurons encode signals that help the animal assess potential risks in the environment, enabling adaptive responses. We plan to investigate dopamine signals in the tail of striatum and other cortical and subcortical areas using our experimental protocol as it is well-suited for examining the temporal derivative nature of dopamine activities. These will help explain the specific mechanisms different brain areas are involved in for learning in aversive situations (Duvarci, 2024).

In real-world scenarios, organisms adaptively adjust their behavior to seek rewards and avoid punishments. This study presents a unified framework explaining how animals navigate complex environments to achieve these goals. Dysfunction in these learning mechanisms appears to be linked to disorders related to aversive situations, such as post-traumatic stress disorder (PTSD) or depression. Beyond enhancing our understanding of everyday decision-making, our results offer valuable insights on developing treatments for these disorders.

## Acknowledgements

This work was supported by IBS-R015-D1, IBS-R015-D2, and National Research Foundation (NRF) of Korea (No. 1711198566 and 2710081229).

## Author contributions

M.P. and H.R.K. conceived and designed the study. M. P. performed most of the experiments, acquired data. and analyzed the data. J.W. performed the initial model-fit analysis and assisted M.P. in further model-fit analyses. H.C. developed the initial version of head-fix avoidance protocol and collected the initial dataset (2 mice). M.P. wrote the first draft manuscript. H.R.K. reviewed and edited the manuscript. H.R.K. supervised the study.

## Data and materials availability

The code written for this study and data will be publicly shared upon peer-review publication.

## Methods

### Mice

A total of 14 adult mice were used in the experiments. All mice were adult C57BL/6J wild-type mice and were used for behavioral experiments with neural recording using dopamine sensor (DA2m). Animals were housed on a 12h dark / 12h light cycle (dark from 07:00 to 19:00) with standard mouse chow and water provided, unless otherwise noted (see Reward-after-shock experiment below). Adult mice (at least 8 weeks old) were used for experiments. All experimental protocols were approved by the Sungkyunkwan University Institutional Animal Care and Use Committee (SKKU IACUC-2024-07-05-1).

### Surgery process

Surgical procedures were under anesthetized conditions induced by 4% isoflurane (0.5-1.0 % at 0.5-1.0 L/min). A mixture of ketamine (0.1 ml/g) and xylazine (10 mg/kg) was injected through intraperitoneal injection before mice were fixed on the stereotaxic frame (Narishige SR-AR). Body temperature was maintained by a heating pad tuned at 37℃. The eyes were covered by eye-ointment after head was fixation via ear-bar on the stereotaxic frame. A skin incision was performed to make a craniotomy at targeted coordinates. The virus (AAV-hsyn-DA2m, 300-400 nL) was loaded and injected into the ventral striatum (bregma 1.1 mm, lateral 1.0 mm, and depths 4.05 - 4.1 mm) using glass micropipette. The injection was performed using a micromanipulator (Nanoliter 2010, WPI) at an estimated rate of 0.01 ul/min for minimal cell damage. The glass pipette was removed slowly after the injection. A head-plate was installed on the skull by applying dental cement on the skull. A mono fiber-optic cannula (200 um diameter, 5mm length, Doric Lenses) was implanted at bregma 1.1, lateral 1.0, and depth 4.1mm and dental cement was applied for fixation. The mice were in recovery for 2-3 weeks after the surgery with ad libitum access to food and water.

### Experimental setup

We conducted experiments in a head-fix virtual reality setup consisting of three LCD monitors (width 50.8 cm, height 30.5 cm) surrounding the animal (Kim et al., 2020). An animal was head-fixed and positioned on the cylindrical treadmill (diameter 20.3 cm, width 10.4 cm). We adjusted the position of the treadmill such that the animals’ eyes were at the center of three monitors. We used ViRMEn software (Aronov & Tank, 2014) to organize the virtual environment and render visual scenes on the monitor. The rotational velocity of the treadmill was measured using a rotary encoder (H1 ball bearing optical shaft encoder, US Digital). Electric shock was delivered through the tail using custom-made coils (Heat-resistant insulated wire stripped, 0.51mm diameter of wire). A pair of coils were tightened on the tail and each coil was connected to the electrical stimulator (A365 Stimulus Isolator, World Precision Instruments).

### Virtual Linear track experiments

#### Standard condition experiment (Track 1)

A virtual linear track was divided into three zones - beginning zone (0 – 60 a.u.), shock zone (60-160 a.u.), and safe zone (160 a.u. - end). Zones were distinguished by different wall patterns (Fig. 1A). Five green donut-shaped circles were used as the wall pattern of the shock zone on both sides. Other zones showed irregular black dots on a navy-blue background. The mouse would be shocked when it stands still in the shock zone but could avoid it by running over 4 a.u./s (corresponds to 0.95 cm/s). We applied electric pulses (amplitude: 0.8 mA, duration: 150 ms) with an interval of 3.5 s. The shock was controlled by analog output from the visual stimulation computer. Because the animal did not voluntarily walk into the shock zone, we adapted the following semi-closed loop condition in the beginning zone: the experiment initially ran in a closed-loop manner. After 6s, the visual scene moved passively at a minimum speed of 2 a.u./s unless the animal ran faster than this. Once the animal reaches the shock zone, the experiment gets back to the complete closed-loop. The scene was turned off during inter-trial interval (ITI) 15 s after mice entered the safe zone. The ITI interval was randomly drawn from an exponential distribution with a minimum of 3.5 s and a maximum of 10 s. After mice were trained in the standard experiment for 5-7 days and mice successfully avoided the shock zone without receiving any shock in over 40% of trials (median success rate of the last training day: 27.9 %, Supp Fig. 1A), mice moved on to the experiment with probe trials added.

#### Speed gain manipulation experiment (Track 1)

Visual gain defines the rate by which the physical running speed of animal (cm/s) is converted to the speed (a.u./s) in the virtual environment. In addition to the standard speed gain (X 1.0), we added speed gains of X 0.5 (scene moved slowly) and X 2.0 (scene moved fast). Slow, standard, and fast gains were used in the 1:3:1 ratio of trials, respectively. The three conditions were randomly interleaved. The minimum scene speed in the beginning zone also increased with the speed gain (2 a.u./s, 4 a.u./s, and 8 a.u./s for the X0.5, X 1.0, and X2.0 conditions, respectively).

#### Teleport experiment (Track 2)

In the teleport experiment, mice were teleported forward or backward to skip or revisit the shock zone. Since the track for the standard experiment (Track 1) has the same pattern in the beginning zone and safe zone, mice could be confused between the two zones when they were teleported. Therefore, for better recognition of the position, we used a different wall pattern for the beginning zone in this experiment (Fig. 3A). The mice were trained without teleportation for 1-2 days in the new track (standard condition only). Once we observed reliable anticipatory running behaviors, we added forward teleport and backward teleport conditions. In the forward teleport condition, the animal was teleported from 57 a.u. to 165 a.u. (the shock zone is 60 a.u. to 160 a.u.). In the backward teleport condition, the animal first underwent the standard task and then was teleported back to the entrance of the shock zone right after they exited (from 165 a.u. to 57 a.u.). At the time of teleportation, a brief blackout of screen (100 ms) helped animals detect the teleportation event. Conditions were randomly interleaved in ratio of 1:1:3, with the highest portion being the standard condition.

#### Reward-after-shock experiment (Track 1 and 3)

A new linear track (Track 3) was introduced to indicate a rewarding context (thick gray plaid pattern at the beginning and safe zone; cyan colored star pattern at the shock zone). In Track 3, a water reward (3.5μL) was delivered through a water port placed in front of the animal’s mouth 1.5s after mice excited the shock zone. The standard condition (Track 1) and reward condition (Track 3) were randomly interleaved with equal probabilities.

### Summary of experimental animals

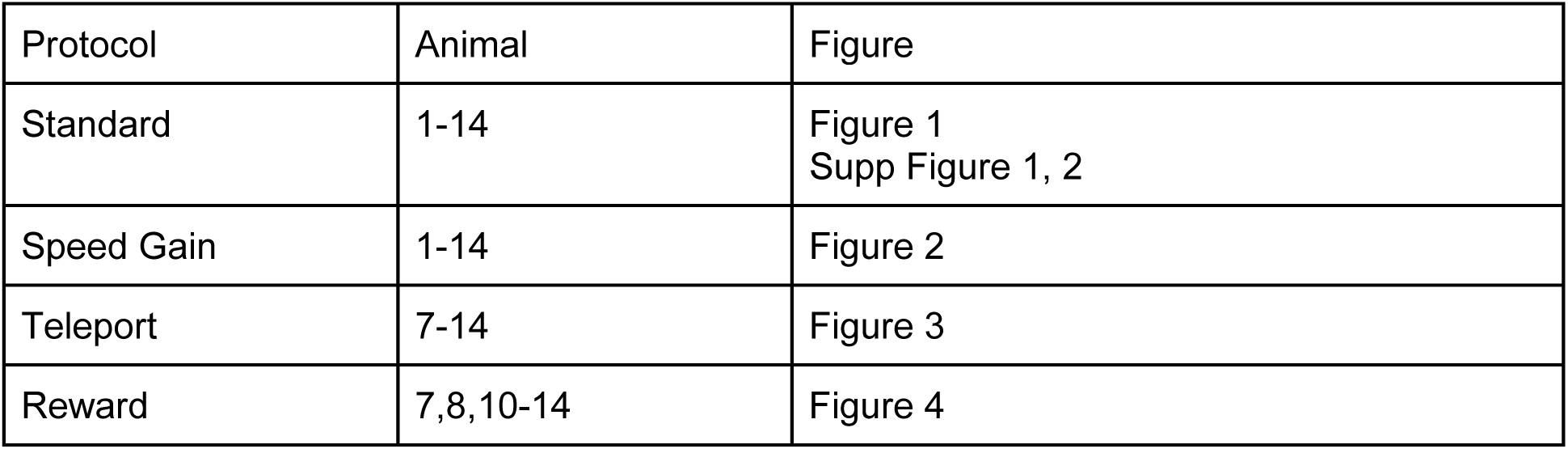

| Protocol | Animal | Figure |
| --- | --- | --- |
| Standard | 1-14 | Figure 1<br>Supp Figure 1, 2 |
| Speed Gain | 1-14 | Figure 2 |
| Teleport | 7-14 | Figure 3 |
| Reward | 7,8,10-14 | Figure 4 |

#### Fiber Photometry and preprocessing

We used fiber photometry to record dopamine activity during the experiments. The AAV vectors that carry G-protein coupled Receptor Activation Based (GRAB) sensor genes (DA2m) were injected into the target brain region (Sun et al., 2020). When GRAB sensors bind with the dopamine, their fluorescence intensity increases. We used a custom-made photometry system to measure the light intensity. It consists of patch cord (400um, Doric Lenses), dichroic mirror (T556lpxr, Chroma), a bandpass filter (ET500/50, Chroma), and photodetector (FDS 100, Thorlabs). The blue laser (473 nm Pulsed Solid-State Lasers, Laserglow Technologies) generated excitation light, which was delivered to the brain by the patch cord. The light went through an optical fiber to the target in the brain. The fiber then collected the emitted fluorescence signals and delivered them to the photodetector. A current preamplifier (SR570, Stanford Research Systems) amplified the signal from photodetector and the voltage signal was digitized by a data acquisition board (NI USB-6002, National Instruments) at 1kHz and saved to the computer using custom MATLAB software.

Recorded voltage signals were preprocessed to obtain z-scored dopamine signals as follows (Kim et al., 2020). The noise from the power line (60Hz) from the raw signal was removed by using a notch filter (signal processing toolbox, MATLAB). The baseline was defined as the lowest 10% of signals using a moving 2-min window. The baseline was subtracted from the raw voltage signal. The results were then z-scored using session-level mean and standard deviation.

### Histology

After completing the task portion of the experiment, animals were perfused using phosphate buffered saline (PBS) followed by 4% paraformaldehyde in PBS. The brain was removed and submerged in 4% paraformaldehyde in PBS for a day. Then, the brain was then moved into a sucrose solution for 2-3 days. Afterwards, the brain was frozen and cut into 100 um slices by cryostat (cryostat model). The brain slices with fiber tracks were loaded on glass slides and stained using 4’,6-diamidino-2-phenylindole (DAPI, Vectashield). Fiber location was determined and recorded referring to standard mouse brain atlas (Paxinos & Franklin, 2019).

### Quantification and statistical analysis

#### Time course analysis

Locomotion (speed), lick, and z-scored dopamine signal were divided by trial and aligned by external events, such as the timing of the animal entering or exiting the shock zone. Lick timestamps were converted to lick rate, a 200ms moving average time window. Locomotion and dopamine responses were not smoothed. The mean and s.e.m. of the event-aligned time course signals were computed across trials to visualize event-related average in peri-stimulus time histogram (PSTH). The s.e.m. of data are represented with shading or error bars. For population time courses, the meantime courses of individual sessions were averaged across animals. For the population-averaged results, shading or error bar represents s.e.m. of the individual means. These population-averaged results were used to summarize the behavior and dopamine responses.

Fluorometry recording experiments were performed with 5-7 days of standard condition and each 2-3 sessions for manipulated experimental protocols. We used the last day’s data from each protocol, as this is when they had the most experience with the modulated protocol condition.

#### Statistical analysis

Statistical analyses were performed using a non-parametric test unless noted otherwise. A Wilcoxon signed-rank test was used for paired tests while Wilcoxon rank-sum test was used for unpaired tests. One, two, and three stars indicate P values less than 0.05, 0.01, and 0.005, respectively. For statistical analysis of time course data, designated time windows were used to compute mean responses. Trial-to-trial responses were used to test for significance in single-session data, and mean responses for each condition were used to test for significance across animals.

#### Permutation test

The permutation test is a non-parametric statistical method used to test if an observed effect is statistically significant. We combined the two datasets for each condition into a single dataset. Then randomly shuffled the combined data and divided into two new groups of the same sizes as the original datasets. We made subtraction between these two new groups and this permutation step was repeated 1000 times for building a distribution of random differences under the null hypothesis. We compared the result with the subtraction of observed results to determine if we can reject the null hypothesis and conclude that the observed difference is statistically significant.

#### Model-fit

For each animal, z-scored dopamine signals were down-sampled to 100Hz and were averaged for each condition aligned by shock zone entrance. To combine the dopamine responses across sessions and experiments for each animal, the single-session average responses were serialized across conditions (e.g., ‘slow’, ’standard’, and ‘fast’ conditions in the speed manipulation experiment). Then the serialized responses were concatenated across sessions (e.g., speed gain experiment and teleport experiments). The concatenation between conditions enabled us to perform fitting using every experimental condition from an individual animal, which can recover the individual value function of an animal. The fitting process was also done by individual sessions without concatenation to compare the model fit result between probe experiments (e.g. speed gain experiment and teleport experiment, Fig. 5C). To address the slow dynamics of DA2m response relative to the latent variable (e.g., TD error), we convolved the fitted latent responses using a kernel. The kernel for DA2m was made with averaged unexpected reward responses from all subjects.

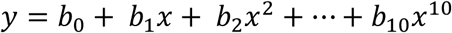

The value function was defined as the 10^th^ polynomial function to reflect the potentially non-monotonic shape of the value function. We used polynomial regression to fit the recorded data (‘regress’ function in MATLAB). Since the deterministic regression function did not allow us to optimize gamma parameter, we used a large set of gamma values ranging from 0.97 to 1.0 in 0.0001 step. We performed polynomial regression for every gamma parameter and chose gamma giving the maximum R^2^ value.

## Figures and figure legends

**Supplementary Figure 1.**
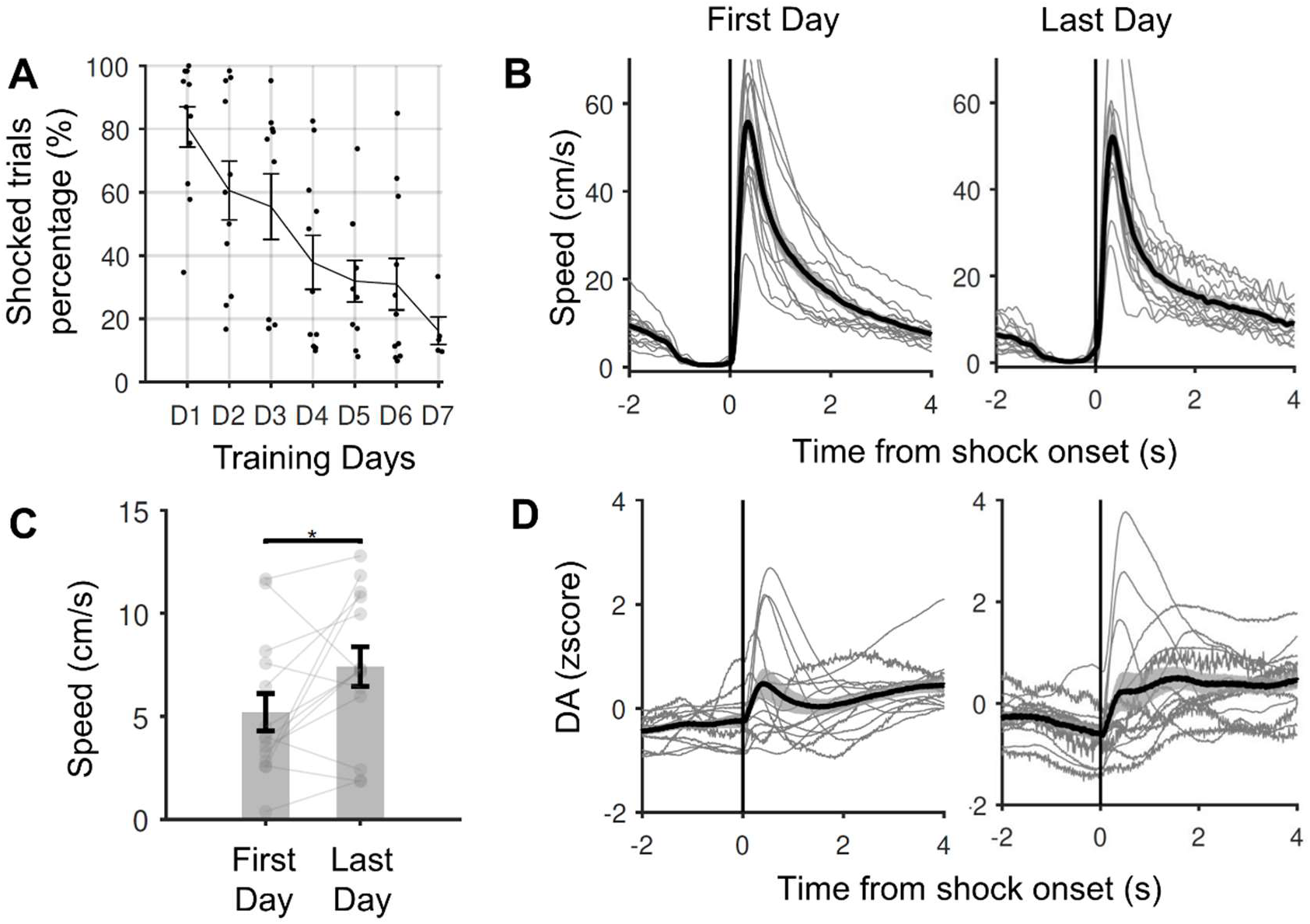
Avoidance learning curve and response to shocks. (A) Percentage of shocked trials by training days of the standard condition (n = 14 mice, t-test, R = -0.97, P = 0.0002). (B) Population of average speed aligned by the time of shock onset in the first day (left) and last day (right) of training standard condition (n = 14 mice). (C) Comparison of speeds in [-3s 0s] before shock zone entrance between the first day and the last day of training (P = 0.041, signed rank test, n = 14 mice). Gray lines indicate individual sessions. (D) Population of averaged dopamine signal aligned by the time of shock onset in first day (left) and last day (right) of training standard condition (n = 14 mice).

**Supplementary Figure 2.**
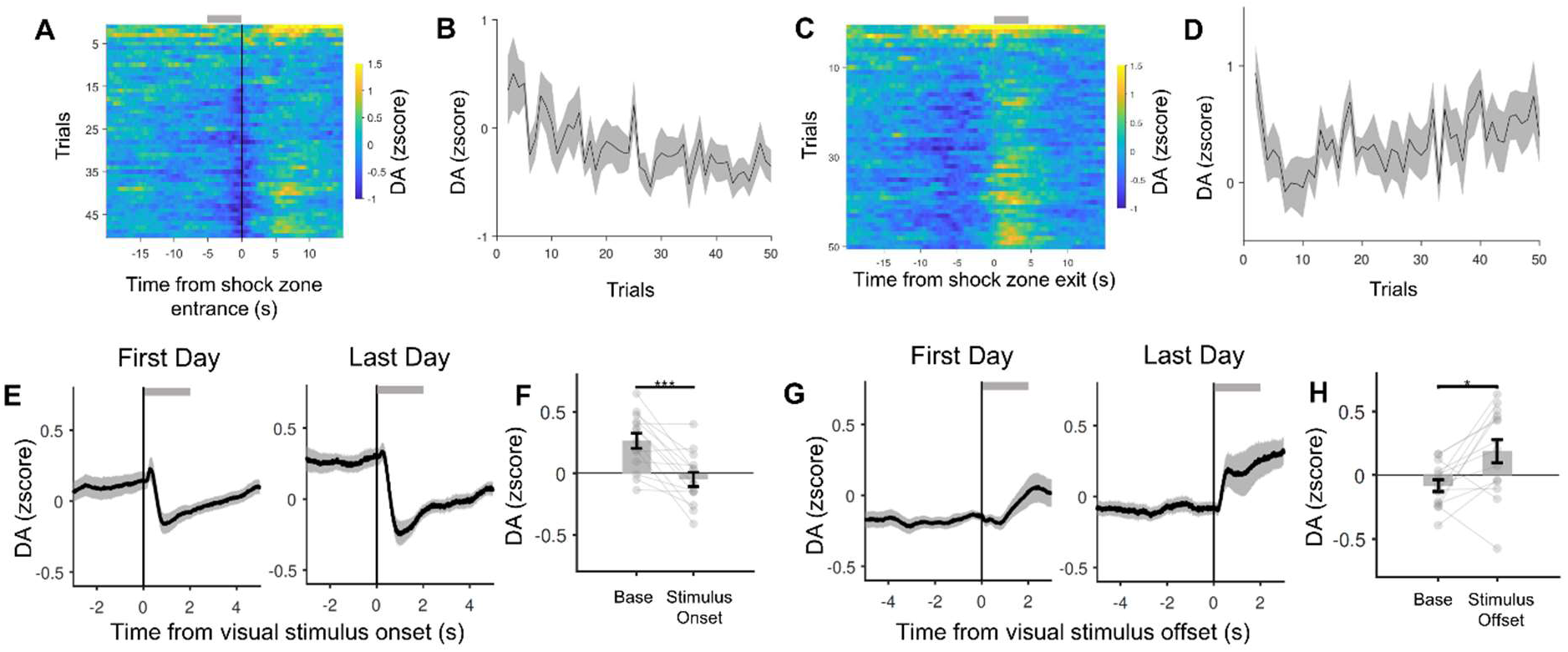
Within- and across-day changes of dopamine activity over training. (A) Heatmap plot of averaged population result trial-by-trial aligned by shock zone entrance at the first day of training (n = 14 mice). (B) Quantified result of dopamine response in time window [-5s 0s] from shock zone entrance averaged for each trial (R = - 0.23, P < 0. 0001, t-test, n = 14 mice). (C) Heatmap plot of averaged population result trial-by-trial aligned by shock zone exit at the first day of training (n = 14 mice). (D) Quantified result of dopamine response in time window [0s 5s] from shock zone exit averaged for each trial (R = 0.11, P =0.003, t-test, n = 14 mice). (E) Dopamine signal of a population result of first day (left, n = 14 mice) and last day (right, n = 14 mice) aligned by the time of visual stimulus onset. (F) Comparison of dopamine signal at the last day of standard experiment in time window of [-3s 0s], [0s 3s] from visual stimulus onset (P = 0.0007, signed-rank test, n = 14 mice). (G) Dopamine signal of a population result of first day (left, n = 14 mice) and last day (right, n = 14 mice) aligned by the time of visual stimulus offset. (H) Comparison of dopamine signal at the last day of standard experiment in time window of [-3s 0s], [0s 3s] from visual stimulus offset (P = 0.012, signed-rank test, n = 14 mice).

**Supplementary Figure 3.**
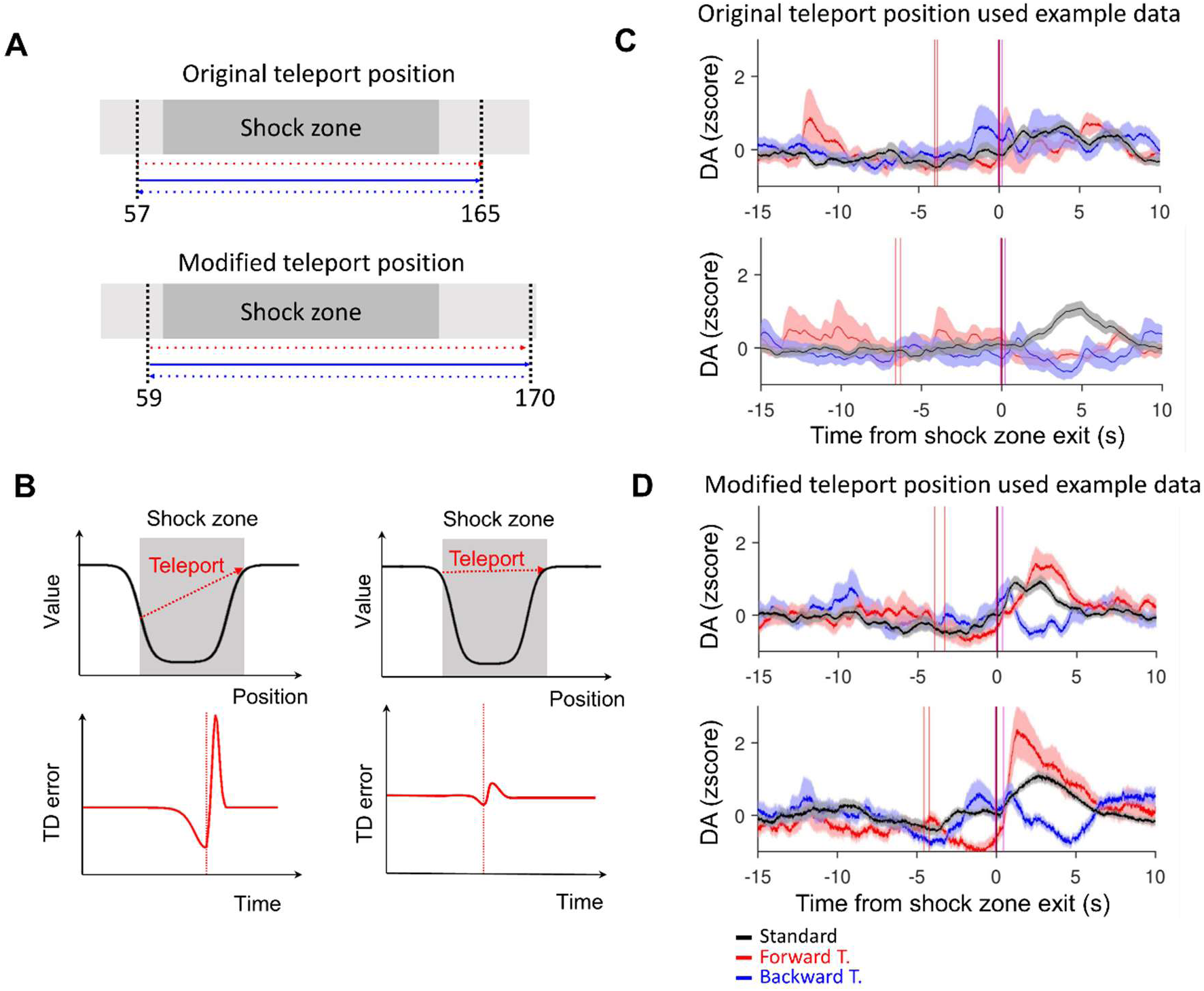
Changing teleport position in teleport experiment. (A) Original and modified teleport position in virtual linear track. (B) Schematic value function and the expected TD error when the position is teleported from the start the end of the shock zone. (C) Dopamine signal of the example session which was recorded using original teleport position (red line, shock zone entrance). (D) Dopamine signal of the example session which was recorded using modified teleport position (red line, shock zone entrance).

